# HDAC6/aggresome processing pathway importance for inflammasome formation is context dependent

**DOI:** 10.1101/2023.08.15.553363

**Authors:** Longlong Wang, Adeline Unterreiner, Ronan Kapetanovic, Selma Aslani, Yuan Xiong, Katherine A. Donovan, Christopher J. Farady, Eric S. Fischer, Frédéric Bornancin, Patrick Matthias

**Affiliations:** Friedrich Miescher Institute for Biomedical Research, 4058 Basel, Switzerland; Faculty of Sciences, University of Basel, 4031 Basel, Switzerland; Novartis Institutes for Biomedical Research, 4056 Basel Switzerland; Department of Cancer Biology, Dana Farber Cancer Institute, Boston, MA 02215, United States; Department of Biological Chemistry and Molecular Pharmacology, Harvard Medical School, Boston, MA 02115, United States

**Keywords:** Inflammasome, HDAC6, degrader, inhibitors, DARPin, interleukin

## Abstract

The inflammasome is a large multiprotein complex that assembles in the cell cytoplasm in response to stress or pathogenic infection. Its primary function is to defend the cell and promote the secretion of pro-inflammatory cytokines, including IL-1β and IL-18. It was shown that in immortalized bone marrow derived macrophages (iBMDMs) inflammasome assembly is dependent on the deacetylase HDAC6 and the aggresome processing pathway (APP), a cellular pathway involved in the disposal of misfolded proteins. Here we used primary BMDMs from mice in which HDAC6 is ablated or impaired and found that inflammasome activation was largely normal. We also used human peripheral blood mononuclear cells and monocytes cell lines expressing a synthetic protein blocking HDAC6-ubiquitin interaction and impairing the APP and found that inflammasome activation was moderately affected. Finally, we used a novel HDAC6 degrader and showed that inflammasome activation was partially impaired in human macrophage cell lines with depleted HDAC6. Our results therefore show that HDAC6 importance in inflammasome activation is context dependent.

## Introduction

The inflammasome is a critical protein complex involved in the body’s immune response (1). When activated, the inflammasome releases the pro-inflammatory cytokines interleukin (IL)-1β and IL-18, which can result in tissue damage and cell death through pyroptosis (2). Several sensor proteins can nucleate inflammasome assembly, including NLRP and NLRC subfamilies, AIM2, and pyrin. Among these, the NLRP3 inflammasome, which is key for sterile inflammation has been extensively studied (3). Activation of the NLRP3 inflammasome often requires a transcriptional priming step to upregulate the inflammasome components such as NLRP3 and IL-1β. Priming agents, e.g., bacterial lipopolysaccharide (LPS), tumor necrosis factor (TNF) or phorbol 12-myristate 13-acetate (PMA), also trigger critical post-translational modifications that sensitize the inflammasome components to subsequent activation. Various stimuli, including the pore-forming toxin nigericin, adenosine-triphosphate (ATP) (4) and monosodium urate crystals (MSU) (5) can stimulate NLRP3 inflammasome assembly, leading to pro-caspase 1 activation, followed by cleavage and release of IL-1β (3). Aberrant inflammasome activity has been linked to many diseases (3), and a detailed understanding of the inflammasome’s activation process is crucial for the development of potential treatments for diseases related to inflammation and immune responses.

The aggresome processing pathway (APP) is a stress-response pathway that is activated when misfolded proteins accumulate; it promotes the formation of the aggresome, a deposit of misfolded and ubiquitinated proteins near the microtubule-organizing center (MTOC), which is subsequently eliminated by autophagy (6). An essential component of the APP is the lysine deacetylase HDAC6 (7), an atypical enzyme with two catalytic domains and a zinc finger domain (ZnF-UbP, hereafter ZnF) binding the small protein ubiquitin (Ub). HDAC6 has been implicated in diverse normal or pathological cellular processes, such as stress response (e.g. APP and stress granules formation) (8), cancer (9), inflammation and viral infection (10). Depending on the situation, either the catalytic domains (regulating acetylation levels of the relevant substrate) and/or the ZnF domain (recruiting Ub) are important for optimal HDAC6 function. In its role in the APP, HDAC6 acts as an adapter protein that promotes transport of ubiquitinated misfolded proteins by dynein motor proteins along the MTs towards the MTOC. For this, both the ZnF and the catalytic domains are critical.

It has recently been proposed that formation of the NLRP3 and pyrin inflammasomes takes place near the MTOC and uses components of the APP machinery and that consequently inflammasome formation is impaired in cells lacking HDAC6 or expressing HDAC6 with a mutated ZnF domain (11). Furthermore, the lysosomal Ragulator complex was recently reported to activate the NLRP3 inflammasome in vivo via HDAC6 (12). Given the apparent similarity between inflammasome formation and APP – in both cases assembly of a complex near the MTOC – it is plausible that HDAC6 could be involved in both pathways, but to define the absolute requirement of HDAC6 in inflammasome activation, a robust profiling of orthogonal methods needs to be tested in different settings. Here we have examined the importance of HDAC6 for inflammasome activation in different cellular systems, including primary mouse cells (BMDMs), peripheral blood mononuclear cells (PMBCs) and human cell lines and manipulated HDAC6 levels through different approaches. Under the conditions tested, we found that the activation of NLRP3 was only partially dependent on HDAC6.

## Results

### NLRP3 inflammasome is activated efficiently in HDAC6 KO and ZnF^m^ BMDMs

Previous work identified the role for HDAC6 in inflammasome activation in immortalized bone marrow-derived macrophages (iBMDMs) (11). To avoid alterations in transcriptome and other cellular processes associated with establishment of an immortalized cell line, we first used primary BMDMs from wild-type (WT) mice as well as two mouse lines in which HDAC6 is either ablated (KO) (13) or contains a W1116A point mutation in the ZnF domain (ZnF^m^, manuscript in preparation), which abolishes ubiquitin binding (10). BMDMs of the different genotypes were first primed by LPS and then challenged by nigericin, which activates the NLRP3 inflammasome by raising intracellular potassium (K^+^) efflux (14) or MSU, which leads to generation of reactive oxygen species (ROS) through activation of NADPH oxidases (5). Inflammasome activation was monitored by examining the cleaved Caspase-1 p20 fragment by immunoblotting and by measuring release of the cytokine IL-1β in the culture supernatant (Fig. 1A). As shown in Figure 1B and 1C, generation of Caspase-1 p20 and production of IL-1β were similar in WT, KO and ZnF^m^ cells. The inflammasome dependence of the IL-1β release was confirmed by using the specific NLRP3 inhibitor MCC950 (16). Because strong inflammasome activation eventually leads to cell membrane damage and release of cellular contents, we examined cellular toxicity by measuring the level of lactate dehydrogenase (LDH) release from cells. Also in this case, the differences between the genotypes were minimal and did not reach statistical significance (Fig. 1D); thus, deletion or mutation of HDAC6 in primary BMDMs does not appear to protect cells from cell death. Collectively, our results in primary BMDMs did not demonstrate a clear dependency on HDAC6 or its ZnF domain for NLRP3 inflammasome activation.

**Figure 1:**
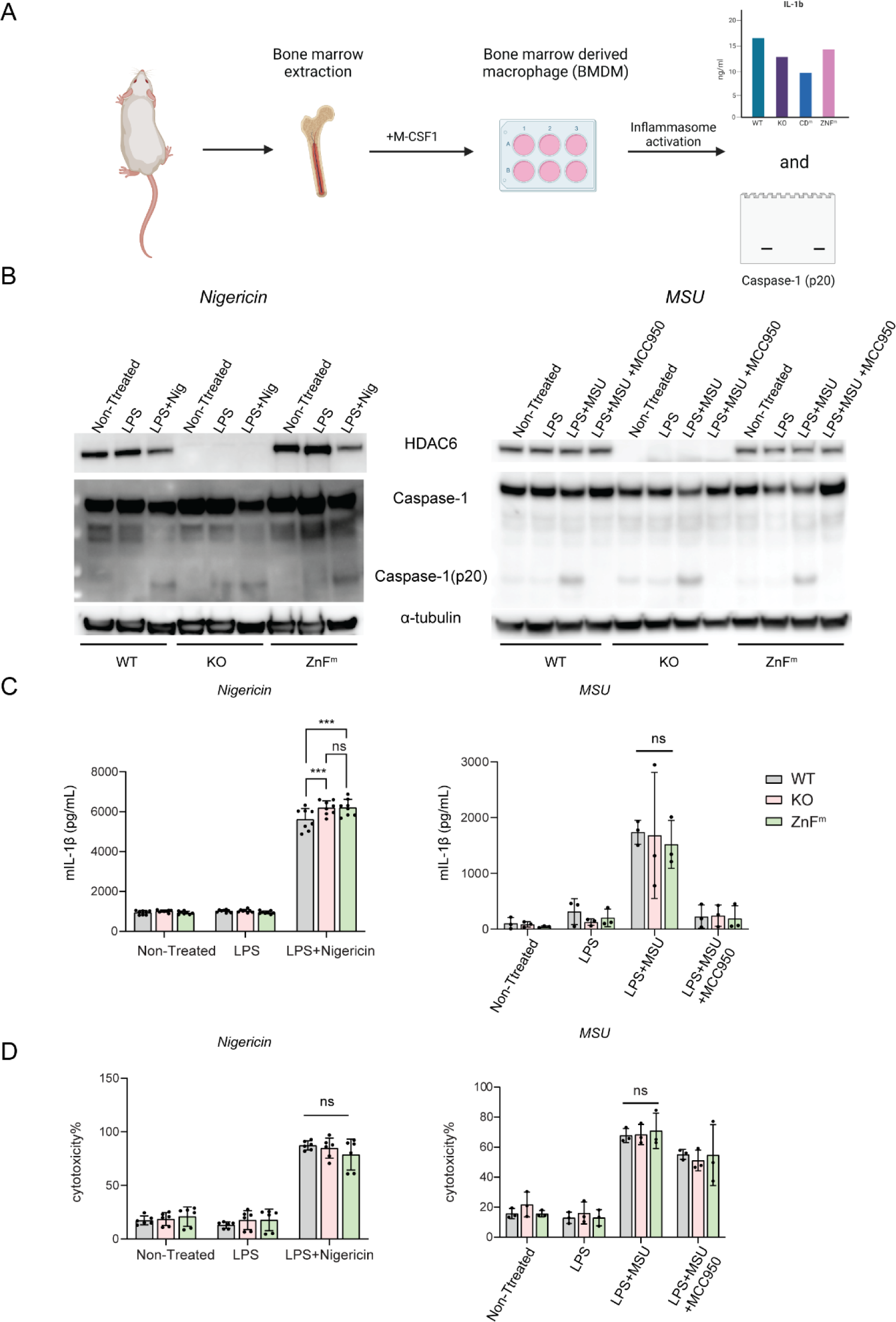
The NLRP3 inflammasome is activated normally in HDAC6 KO or ZnF^m^ primary mouse BMDMs. (A) Workflow for evaluating NLRP3 inflammasome activation in BMDMs. Bone marrow was isolated from mice strains with different mutations in HDAC6 as indicated and BMDMs were expanded for 7 days in the presence of M-CSF1. Subsequently, BMDMs were primed for 4 hrs with LPS and NLRP3 inflammasome formation was induced by nigericin or MSU addition. Release of IL-1β was monitored by ELISA and activation/cleavage of Caspase-1 was evaluated by immunoblotting. (B) Normal NLRP3 activation in WT, HDAC6 KO and HDAC6 ZnF^m^ BMDMs. Immunoblot monitoring HDAC6, Capase-1 (p20) cleavage and α-tubulin as loading control. nigericin and MSU were used with LPS primed BMDMs (n = 3). (C) Normal mIL-1β release in the supernatant from WT, HDAC6 KO and ZnF^m^ BMDMs. IL-1β from (B) was measured by ELISA and statistical analysis was done with one-way ANOVA. ns, no significant difference. (D) No difference in cytotoxicity following inflammasome induction in WT, HDAC6 KO and ZnF^m^ BMDMs. Lactate dehydrogenase (LDH) was measured in the culture supernatant from (B) and statistical analysis was done by one-way ANOVA. ns, no significant difference.

### A ZnF-specific DARPin moderately inhibits NLRP3 inflammasome activation in THP-1 cells

To investigate more broadly the role of the HDAC6 ZnF in NLRP3 inflammasome activation, we switched from primary BMDMs to the leukemic THP-1 monocyte/macrophage cell line, a widely used surrogate for human monocytes or macrophages. We recently reported a synthetic protein based on a designed ankyrin repeat protein (DARPin) scaffold, DARPin F10, which can selectively bind to the HDAC6 ZnF Ub binding pocket in cells and blocks Ub recruitment (15) (Fig. 2A). We previously made a degradable version of DARPin F10 by fusing it to the FKBP^F36V^ degron tag (16) and showed that it is efficiently degraded in A549 cells upon addition of the chemical dTAG-13 (hereafter dTAG). We used this system to generate THP-1 cells expressing DARPin F10 (F10-FKBP^F36V^ THP-1 cells) and found that the protein was well-expressed and could be efficiently degraded within 6 hours of dTAG addition (Fig. 2B). Nigericin-induced NLRP3 activation was comparable in dTAG treated parental (WT) THP-1 cells (Fig. 2C) and in control cells expressing only the FKBP^F36V^ moiety (FKBP^F36V^ THP-1 cells, Fig. S1), thus excluding a possible inhibitory effect of FKBP^F36V^ expression or dTAG on IL-1β release. When parental WT THP-1 cells were treated with MCC950 (17), or CGP084892, a Caspase-1 inhibitor (18), IL-1β release was strongly reduced, as expected. A modest reduction of IL-1β secretion was observed in F10-FKBP^F36V^ THP-1 cells, when compared to the WT THP-1 cells. Degradation of DARPin F10 by dTAG treatment led to a partial, but significant, recovery of IL-1β release (Fig. 2C). We next examined early timepoints - 0.5 and 1.5 hr post-nigericin activation - to study the impact of DARPin F10 on the kinetic of IL-1β release. In agreement with Figure 2C, we observed again a slight increase of IL-1β at 3.5 hr in dTAG-treated cells (Fig. 2D). However, a larger and significant difference was seen at 1.5 hr post activation. These data indicate that blockade of the HDAC6 ZnF domain moderately inhibits the NLRP3 inflammasome, particularly during the early phase of activation by nigericin in THP-1 cells.

**Figure 2:**
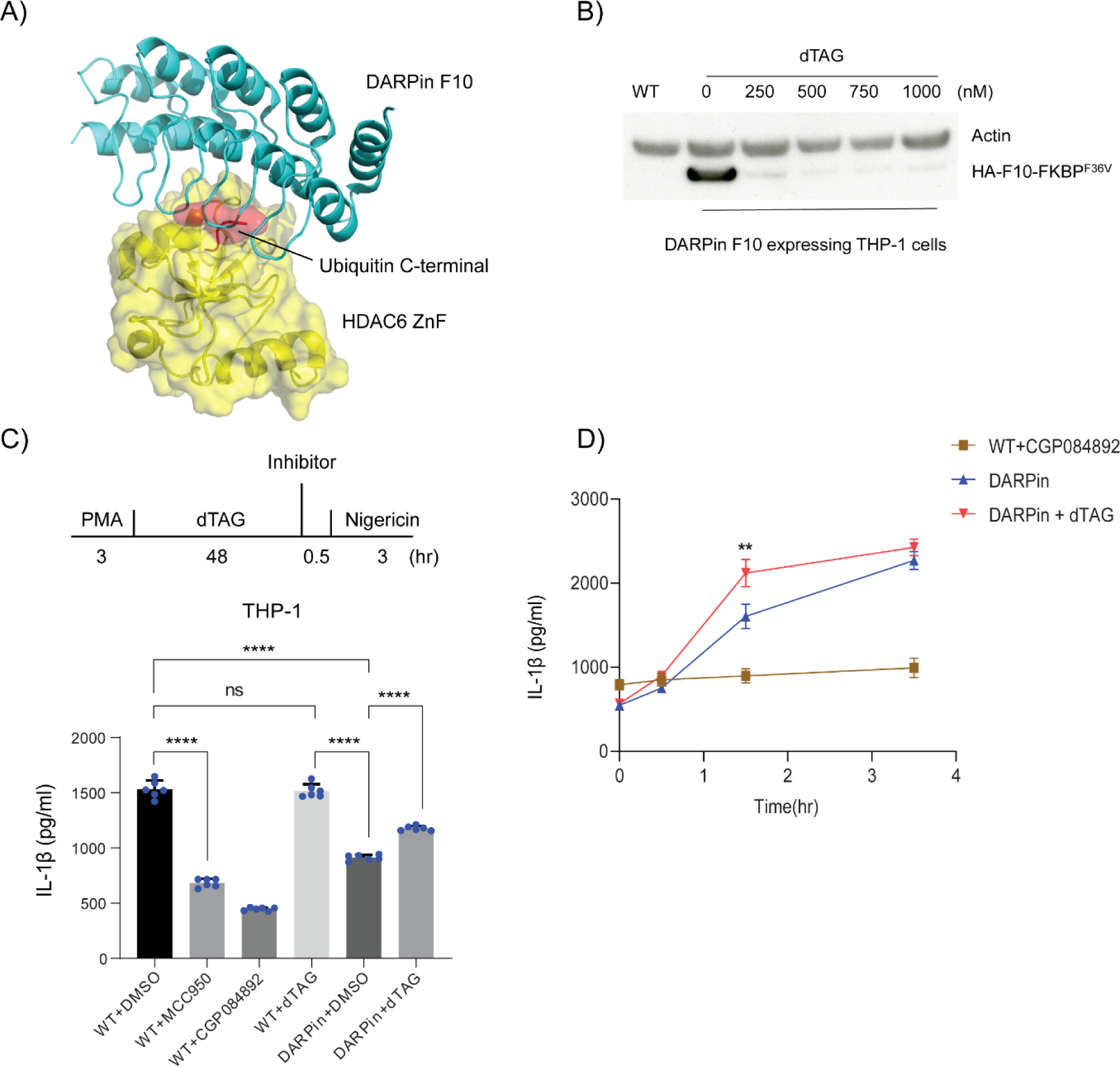
A DARPin blocking the HDAC6-Ub interaction has a modest effect on IL-1β release from THP-1 cells. (A) Structure of the DARPin F10 - HDAC6 ZnF domain complex (PDB: 7ZYU) showing blockade of Ub recruitment. The surface representation shows HDAC6 ZnF (yellow) and the C-terminal LRGG of Ub (light pink) (overlayed with PDB:3GV4), the ribbon representation depicts DARPIN F10 (cyan). (B) Establishment of a THP-1 F10-FKBP cell line with degradable DARPin F10. Immunoblotting with lysates of THP-1 F10-FKBP cells treated at the indicated concentration with dTAG (for 6 hrs). The leftmost lane (WT) shows the parental THP-1 cells. The membrane was probed with antibodies against actin as loading control and F10 (HA-F10-FKBP ^F36V^, detected with anti-HA). (C) IL-1β release under various treatments in THP-1 F10-FKBP cells at 3 hrs post-nigericin stimulation. Cells were treated as indicated in the scheme at the top. The graph at the bottom shows the measurement of IL-1β in cell supernatants, based on 3 independent experiments. MCC950 and CGP084892 are NLRP3 and Caspase-1 inhibitors, respectively. One-way ANOVA test was applied for statistical analysis. ****, p<0.0001. (D) IL-1β release from 0 hr to 3.5 hr after nigericin treatment in THP-1 or THP-1-F10-FKBP cells. Experimental set-up was as in (B) and IL-1β concentration was monitored at timepoint 0, 0.5, 1.5, and 3.5 hr post nigericin treatment. Two-way ANOVA tests were applied, and comparison was performed within each timepoint between with and without dTAG group. **, p<0.01.

### HDAC6 degradation moderately inhibits NLRP3 inflammasome in PBMCs

Blocking specifically the ZnF domain in THP-1 cells by DARPin F10 showed a partial inhibition of NLRP3 inflammasome activation, confirming that HDAC6 contributes to this pathway. To extend these observations in other cell settings, we next made use of a novel HDAC6 proteolysis targeting chimera (PROTAC) degrader, XY-07-35 (19). Treatment of cells with XY-07-35 leads to recruitment of the E3 ligase cereblon (CRBN) to HDAC6, ubiquitination and subsequent proteasomal degradation of HDAC6 (Fig. 3A). This degrader is based on the HDAC inhibitor suberoylanilide hydroxamic acid (SAHA) fused by a linker to pomalidomide as a CRBN-binding moiety; the negative control compound XY-07-191 contains a methyl group on the pomalidomide moiety which disrupts CRBN recruitment (Fig. 3B). *In vitro* enzymatic assays with purified HDACs showed that XY-07-35 inhibited HDAC6 with a low IC_50_=0.0485 μM, which is 10 times lower than for HDAC8 (0.621 μM) or other HDACs (1.04-5.75 μM; Fig. 3C); in addition, the degradation profile of XY-07-35 is highly specific for HDAC6 (19).

**Figure 3:**
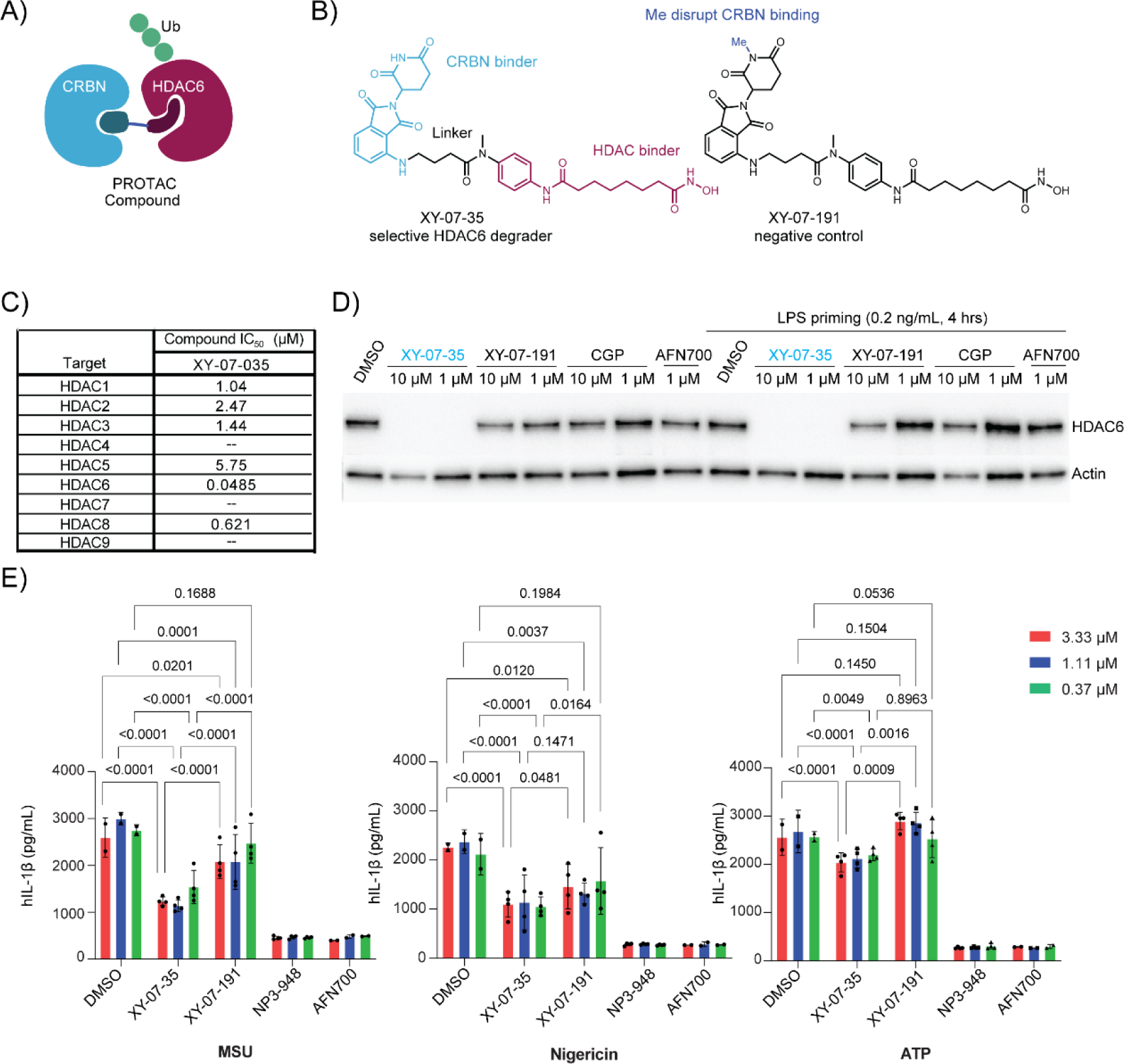
Effect of XY-07-35 mediated HDAC6 degradation on NLRP3 inflammasome activation in PBMCs. (A) Schematic of the mode of action of an HDAC6 degrader. The PROTAC engages both HDAC6 and the E3 ligase cereblon (CRBN), which then leads to ubiquitination and degradation of HDAC6. (B) Chemical structure of the HDAC6 degrader XY-07-35 and of the negative control XY-07-191 which no longer recruits CRBN. (C) *In vitro* enzymatic assay of compound XY-07-35 on the activity of different HDACs. IC^50^ for each enzyme is shown. (D) XY-07-35 mediated HDAC6 degradation in PBMCs. PBMCs (without or with LPS priming for 4 hrs) were treated with the indicated compounds and HDAC6 levels were detected by immunoblotting with an anti-HDAC6 antibody. Detection of actin was used as control for loading. CGP, CGP084892 (Caspase-1 inhibitor); AFN700 is an inhibitor of the NF-kB pathway (IKK inhibitor). (E) HDAC6 degradation impact on IL-1β release in PBMCs following MSU, nigericin or ATP treatment. The scheme at the top depicts the experimental outline. The graphs below show the results of 3 independent experiments using the inducers as indicated. NP3-948 is an inhibitor of the NLRP3 inflammasome.

We first tested the efficacy of XY-07-35 in immune cell lines. Unexpectedly, XY-07-35 treatment did not lead to HDAC6 degradation in both non-differentiated and PMA-differentiated THP-1 cells (Fig. S2). Therefore, to assess its effect on NLRP3 inflammasome activation in human cells we used peripheral blood mononuclear cells (PBMCs). HDAC6 was efficiently degraded in PBMCs at 1 μM XY-07-35 concentration; by contrast, the negative control compound XY-07-191 or the IκKβ inhibitor AFN700 (18) had only minimal or no effect on HDAC6 levels (Fig. 3D). To test the functional consequence of HDAC6 degraders, PBMCs were first treated with the HDAC6 degrader or negative control compound for 24 hrs, followed by LPS priming and subsequent stimulation with MSU, nigericin, or ATP to activate the NLRP3 inflammasome (Fig. 3E). The HDAC6 degrader XY-07-35 clearly reduced the IL-1β release under all stimulatory conditions as compared to DMSO, though partial inhibition was also seen with the negative control XY-07-191, suggesting some non-specific effects of the scaffold. HDAC6 degradation had the most profound inhibition of MSU-triggered IL-1β release, achieving ca. 60% of IL-1β reduction as compared to DMSO, while under nigericin and ATP stimulation, the reduction was 40%. A statistically significant reduction of IL-1β between XY-07-35 and the control compound XY-07-191, was only seen in the MSU conditions. Inhibitors of other nodes of the pathway, the NLRP3 inflammasome inhibitor NP3-948 and AFN700 both led to an almost complete (>90%) inhibition of IL-1β release. To better elucidate the inhibitory effect of XY-07-35 on IL-1β release, we calculated the inhibition ratio of IL-1β release (Fig. S3). The negative control compound XY-07-191 also minimally inhibited IL-1β release, but less effectively than XY-07-35, suggesting a beneficial effect of HDAC6 degradation. We also measured cell viability by using resazurin fluorescence in PBMCs, but since monocytes constitute only a minor fraction of cell population, the pyroptosis could not be tracked efficiently. Overall, these results show partial reduction in NLRP3 inflammasome activation following degradation of HDAC6, with the magnitude of the effect depending on the stimulus.

### HDAC6 degradation does not affect pyrin inflammasome activation

According to a previous report (11), activation of both NLRP3 and pyrin inflammasomes depends on HDAC6. To evaluate the importance of HDAC6 for the pyrin inflammasome, we used a previously established U937 pyrin-overexpressing cell line (20). In these cells, treatment with the bile acid derivative BAA473 activated the pyrin inflammasome, leading to the release of cytokines (e.g., IL-1β, IL-18) and pyroptosis. HDAC6 was efficiently degraded in the U937-pyrin expressing cells by XY-07-35 at concentrations ranging from 0.1 μM to 10 μM (Fig. 4A). However, treatment with XY-07-35 did not result in any change in IL-18 secretion compared to either DMSO control or the XY-07-191 control (Fig. 4B). Additionally, we measured cell viability and found that HDAC6 degradation by XY-07-35 did not protect cells from pyroptosis (Fig. S4), confirming that in this setting, HDAC6 degradation has no effect on pyrin inflammasome activation.

**Figure 4:**
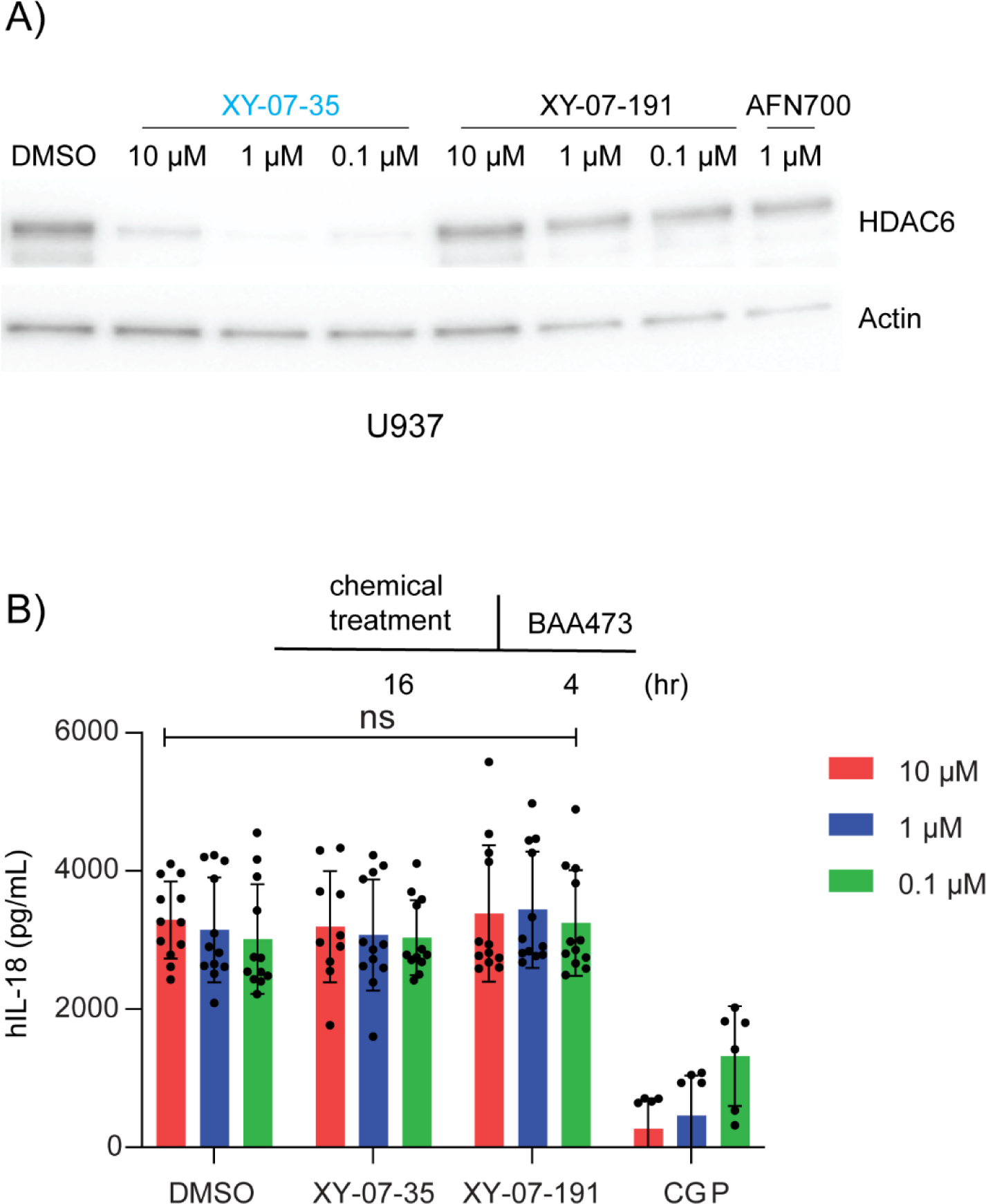
Effect of HDAC6 degradation on pyrin inflammasome activation in U937 cells. (A) XY-07-35 mediated HDAC6 degradation in pyrin-expressing U937 cells. Cells were treated with the indicated compounds and HDAC6 was detected by immunoblotting with an anti-HDAC6 antibody. Detection of actin was used as a control for protein loading. (B) No impact of HDAC6 degradation on IL-18 release in U937-pyrin cells treated with the indicated compounds. The scheme at the top depicts the experimental outline. The graph at the bottom presents IL-1β release based on 4 independent experiments. CGP, CGP084892.

## Discussion

In this study, we utilized a range of cellular systems and several orthogonal approaches to modulate HDAC6 activity and probe its role in NLRP3 and pyrin inflammasome activation. HDAC6 protein levels were modulated by both knock-out and chemically induced degradation, while function was probed by a knock-in unable to bind to ubiquitin and a DARPin inhibitor blocking ubiquitin recruitment. In contrast to a previous study reporting on the HDAC6-ubiquitin interaction as a crucial regulator of inflammasome activation (11), we did not observe a significant difference in NLRP3 inflammasome activation between WT and HDAC6 knock-out or HDAC6 ZnF^m^ (W1116A) BMDM cells. To investigate this further, we examined the effect of a small protein (DARPin F10) binding to the HDAC6 ZnF domain and preventing ubiquitin recruitment. This protein was shown to have a strong inhibitory impact on processes involving the ZnF-Ub interaction, such as the formation of aggresomes or stress granules, or infection by influenza or Zika virus (15). But under conditions of inflammasome induction, we observed only partial downregulation of IL-1β release in THP-1 cells expressing DARPin F10 when stimulated with nigericin. In primary PBMCs and U937 cells, the selective HDAC6 degrader XY-07-35 efficiently degraded HDAC6 but, HDAC6 had only modest, stimulation-dependent effects on NLRP3 inflammasome activation. Furthermore, HDAC6 degradation had no effect on IL-18 release and pyroptosis mediated by pyrin. With respect to NLRP3 inflammasome activation, HDAC6 degradation had the strongest effect on MSU-stimulated IL-1β release, which is a milder, non-pyroptotic form of NLRP3 activation (Fig. S3). Similarly, in THP1 cells, HDAC6 inhibition seemed to have its most pronounced effect at early timepoints. It is possible that HDAC6 plays a more pronounced role in response to crystalline inflammasome activators (in agreement with the results in Magupali et al, 2022). It has also been noted that other NLRP3 regulatory mechanisms such as PKD phosphorylation play an important role at early time points in cell culture experiments, but compensatory pathways override these effects at later timepoints (21).

Based on our results it seems that the importance of HDAC6 for NLRP3 inflammasome activation is not absolute, but condition and context dependent. On one hand, we noted that blocking the HDAC6 ZnF-Ub interaction with DARPin F10 in THP-1 cells and degrading HDAC6 in PBMCs both resulted in an inhibitory effect, thus supporting a role for HDAC6 in this pathway. When degrading DARPin F10 with dTAG we only observed a partial rescue of IL-1β release; this can possibly be explained by incomplete degradation of DARPin F10, leading to some residual F10 (even in tiny amounts) that could interfere with ubiquitin recruitment. Similarly, incomplete HDAC6 clearance by the proteasome system might be responsible for the moderate downregulation of IL-1β release or cell death protection in XY-07-35 treated PBMCs. Furthermore, we found that the negative control compound XY-07-191 also exhibited slight inhibition of NLRP3 inflammasome activation, which we attribute to its possible catalytic inhibition of HDAC6. Indeed, both compounds are derived from SAHA (Fig. 3B), which is known to inhibit all HDACs including HDAC6, and previous studies have demonstrated that HDAC inhibition can downregulate IL-1β release (11, 22).

On the other hand, we found that the activation of the NLRP3 inflammasome in BMDMs from HDAC6 KO and ZnF^m^ (W1116A) mice took place normally; likewise, activation of the pyrin inflammasome in U937-pyrin cells was not affected by HDAC6 depletion. In a previous study using immortalized BMDMs (11), researchers described a HDAC6-mediated aggresome-like pathway for NLRP3 inflammasome formation, and reported that deletion of HDAC6 or mutations in the ZnF domain completely blocked inflammasome activation, in contrast to our results. Likewise, another recent study reported that the lysosomal Ragulator complex is important for NLRP3 inflammasome activation in vivo via interaction with HDAC6 (12).

We hypothesize that the partial discrepancy may result from the differences in the cell types and specific experimental conditions used. Immortalized cells are known to carry chromosomal abnormalities that allow them to proliferate indefinitely, leading to changes in their transcriptome, proteome and reliance on specific pathways compared to primary cells. As a result, it is reasonable to speculate that established iBMDMs may behave differently from primary BMDMs. As for the pyrin inflammasome, we used U937 cells instead of iBMDMs, and bile acid derivatives instead of Clostridioides difficile toxin B (TcdB) to activate the pathway, which may also contribute to the observed differences in results obtained. In summary, we conclude that HDAC6 can participate in NLRP3 inflammasome activation, but its importance appears to be context dependent.

## Experimental procedures

### Cell Culture

THP-1, U937, PBMC and BMDM cells were cultured in Roswell Park Memorial Institute (RPMI) 1640 Medium–stable Glutamine (Bioconcept#1-41F03-1) supplemented with 10% fetal bovine serum (FBS). For BMDMs the medium also contained M-CSF 1 (100 ng/mL). All cells were cultured in 5% CO_2_ at 37°C.

### BMDMs isolation and culture

BMDMs were isolated as previously described (23). In short, mice (8∼14 weeks old) were sacrificed and both legs were sterilized in 70% ethanol. Attached tissues were removed by dissection with scissors and the isolated bones were flushed or crushed twice in a mortar with 5 mL RPMI 1640 medium. The suspension from crushed bones was filtered through 100 μm cell strainer (Falcon#352360) to remove debris. Filtered cells were centrifuged at 500 x g, 15 mins at room temperature. Supernatant was removed and cell pellets were resuspended in RPMI 1640 (+M-CSF 1, 100 ng/mL). The cell suspension (from 1 leg) was divided into four Nunc Square 25-cm dishes (Thermo Scientific#166508), with each plate containing 10 mL cell suspension. On Day 3 post-seeding, 5 mL fresh RPMI 1640 (+M-CSF 1) medium was added to each plate. On Day 6, BMDMs were ready for collection or re-seeding for new experiments. All mice handling was done according to Swiss federal guidelines for animal experimentation and approved by the FMI Animal committee and the local veterinary authorities (Kantonales Veterinäramt of Kanton Basel-Stadt).

### Immunoblotting

Treated BMDM cells were lysed with RIPA buffer (supplied with cOmplete™ Protease Inhibitor Cocktail, Merck#11697498001), and protein concentration was determined by Bradford assay. Same protein amounts were loaded onto 4-12% NuPAGE Gels and protein was transferred to PVDF membranes by using Trans-Blot® Turbo™ Transfer System (Biorad#17001917). Membrane was blocked with 5% milk-TBS, and then incubated with primary antibodies overnight at 4°C. Signal was developed with HRP-conjugated secondary antibodies and Amersham ECL Western Blotting Detection Reagent (Cytiva#RPN2106). Antibody information: mouse Caspase-1 (p20) (AdipoGen#AG-20B-0042-C100), mHDAC6 (homemade), human HDAC6 (CellSignaling#7558), α-tubulin (Sigma-Aldrich #T9026), HA (CellSignaling#3724), pan Actin (Sigma-Aldrich#SAB4502632).

### ELISA (for IL-1β and IL-18)

Supernatant from treated cells were collected and frozen at −80°C before determining cytokine concentrations. mIL-1β was determined by ELISA MAX™ Standard Set Mouse IL-1β kit (BioLegend# 432601). Human IL-1β and IL-18 were determined with Cisbio HTRF Cisbio Human IL1β (#62HIL1BPET) and Human IL18 (#62HIL18PEG) protocols. All the data were from 3 or 4 independent biological replicates.

### LDH measurement

Culture supernatants from the different BMDMs samples were analyzed with the CytoTox 96® Non-Radioactive Cytotoxicity Assay Kit (Promega# G1780). Total cell lysate of non-treated (i.e., DMSO only-no inhibitor-, but induced for inflammasome formation) BMDM cells was defined as 100% cell death or cytotoxicity. Values from other conditions were calculated as follows: (OD_treated cells_-OD_no-inhibitor control_)÷(OD_total lysate_-OD_no-inhibitor control_) * 100.

### Single cell clonal THP-1 F10 and FKBP cells establishment

Lentiviral vectors for HA-DARPin F10-FKBP^F36V^ and HA-FKBP^F36V^ was from previous work (15). Lentivirus was generated with 3^rd^ generation lentivirus packaging systems. In detail, DARPin F10 vector and lentivirus packaging plasmids (Tat, Rev, Gag/pol, Vsv-g) were co-transfected in 293T cells in 6 well-plates. After 3 days culturing, supernatants were harvested and concentrated with Lenti-X™ Concentrator (Takara# 631232). Virus pellet was resuspended in 1∼4 mL RPMI1640 medium (with 10% FBS). Next, THP-1 cells were seeded in 6 well-plates (each well with 1 x10^6^ cells), and then lentivirus was added to the plate in the presence of polybrene (8 μg/mL), followed by centrifugation at 1500 rpm / 453 x g, for 45 mins at room T°. After transduction, THP-1 cells were cultured at 37°C for 7∼14 days. The cell number and density were monitored carefully; when a concentration of > 2 x10^6^ cells per mL was reached, 2 μg/mL puromycin was added for selection for 3∼6 days. Surviving cells were single cell sorted by FACS into 96 well plates (in RPMI1640 without puromycin). After about 1 month in culture, single cell clones were analyzed by immunoblotting to test for DARPin F10 expression. Cell clones with a good expression level were stored in cryo-solution (90% FBS + 10% DMSO) in liquid nitrogen.

### Inflammasome activation

For BMDM cells, 1.5 x10^6^ cells per well were seeded in 6 well-plates. After priming with 1 mg/ml LPS (Invivogen#tlrl-b5lps), cells were challenged with 200 μg/ml MSU crystals (Invivogen#tlrl-msu-25) for 6 hours or 20 μM nigericin (Sigma-Aldrich#N7143-5MG) for 30 mins. For THP-1 cells, 5×10^5^ cells/mL were primed with 0.5 μM PMA (Invivogen#tlrl-pma) in T75 Ultra-low binding flask (CORNING#3814) for 3 hours, and then cells were collected by centrifugation at 500 x g, for 5 mins. The supernatant was removed and fresh medium was added to have cells at a density of around 5×10^5^ cells/mL. 100 μL cells were seeded in 96 well plate and incubated for 1∼2 days. During the culturing, dTAG was added when needed to degrade the DARPin. After 1∼2 days culturing, inflammasome was activated by adding 15 μM nigericin for up to 4 hours. For PBMCs, 3×10^4^ cells per were seeded, and primed with 0.2 ng/mL LPS for 4 hours and then incubated with MSU 200 μg/mL (5 hours), nigericin 2 μM (2 hours) or ATP 3 mM (0.5 hours). For pyrin inflammasome, U937-pyrin cells were seeded in 96 well after 0.5 μM PMA treatment for 3 hours and then challenged by BAA473 (200 μM) for 4 hrs at 37°C. For control experiments inhibitors were used at the following final concentration: MCC950 at 10 μM, CGP084892 at 20 μM. NP3-948 is a potent NLRP3 inhibitor developed at Novartis that competes with MCC950 for binding to the NLRP3 protein (24).

### Cell viability Assay

Cell viability of PBMCs and U937 cells was checked with PrestoBlue™ HS Cell Viability Reagent (Invitrogen# P50200) according to the manufacturer’s instructions. After incubation for 1 to 2 hrs, fluorescence was determined with a PheraStar plate reader under Ex/Em=540/590 nm.

### Statistical analysis

All the statistical analysis were done with One-way or Two-way ANOVA test, corrected for multiple comparisons using post-hoc significance Tukey test. A significance level of p < 0.05 was used.

## Data availability

Mass spectrometry data related to HDAC6 degrader XY-07-35 and XY-07-191 are deposited to PRIDE under archive number PXD023652. The data that supports the findings described in this work are available from the corresponding author upon request.

## Acknowledgements and funding

We thank Christelle Arnold (NIBR) for her help in setting up the collaboration with FMI. We also thank Chun Cao and Gabriele Matthias for providing HDAC6 KO and ZnF^m^ mice. Financial support through the SystemsX.ch MRD project VirX 2014/264, evaluated by the Swiss National Science Foundation, and by the European Research Council ERC Synergy grant number 856581 CHUbVi is gratefully acknowledged. This work was supported by the Novartis Research Foundation. This work was supported by the National Institutes of Health (NIH) grant R01CA262188 (to E.S.F.).

## Authors contributions

L.W. performed most of the experiments in BMDMs, THP-1 F10 cells, analysis, preparation of figures and wrote the manuscript draft. R.K helped with BMDMs isolation and culture, and with manuscript modifications. X.Y., K.A.D. and E.S.F. were responsible for providing the HDAC6 degrader and corresponding characterization, and manuscript proofreading. F.B. led the collaboration between NIBR and FMI, and helped with experimental design and manuscript preparation, A.U. was responsible for the experiments in PBMCs and U937-pyrin cells, and for measuring IL-1β in various samples, S.A. performed the HDAC6 degrader experiments in THP-1 and U937 cells, C.J.F. was responsible for experimental design, conceptualization, and manuscript preparation. P.M was responsible for the conceptualization, supervision, writing the original draft and review, project administration and funding acquisition. All authors contributed to the final manuscript.

## Supporting information

**Figure S1 (related to Fig. 2):**
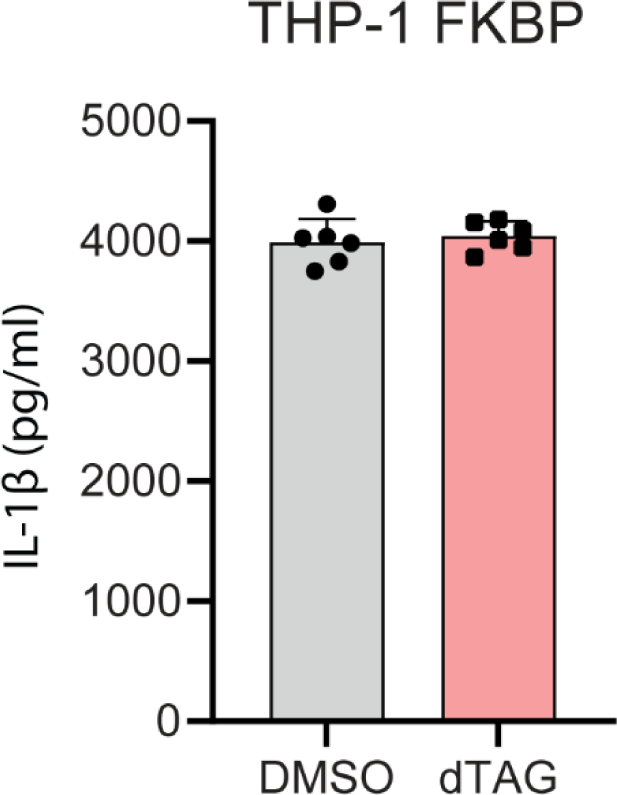
THP-1 FKBP cells were established for evaluating the dTAG effect on IL-1β release. Cells were primed by PMA and then treated with nigericin as in Figure 2.

**Figure S2 (related to Fig. 3):**
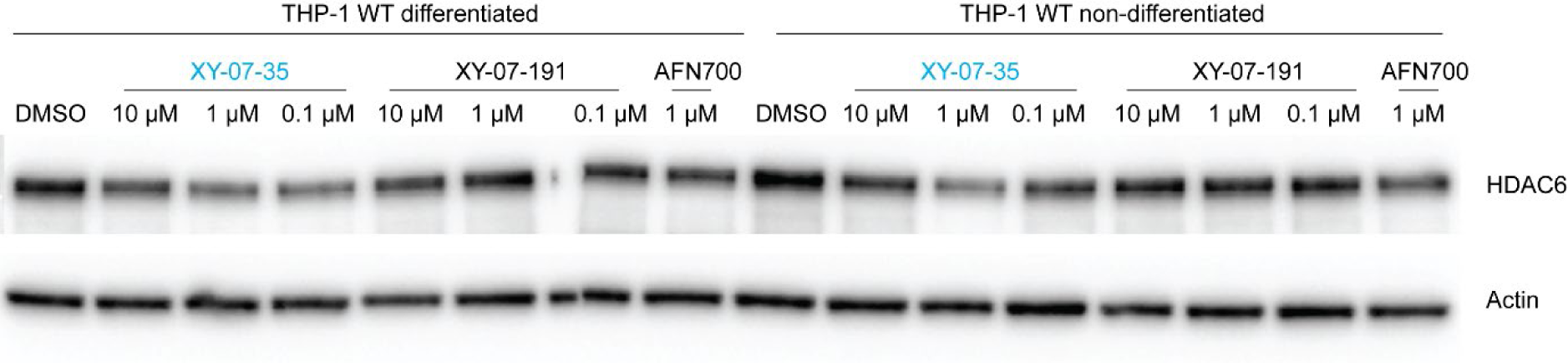
THP-1 cells, with or without in vitro differentiation, were incubated with XY-07-35 and the HDAC6 level was examined by immunoblotting.

**Figure S3 (related to Fig. 3):**
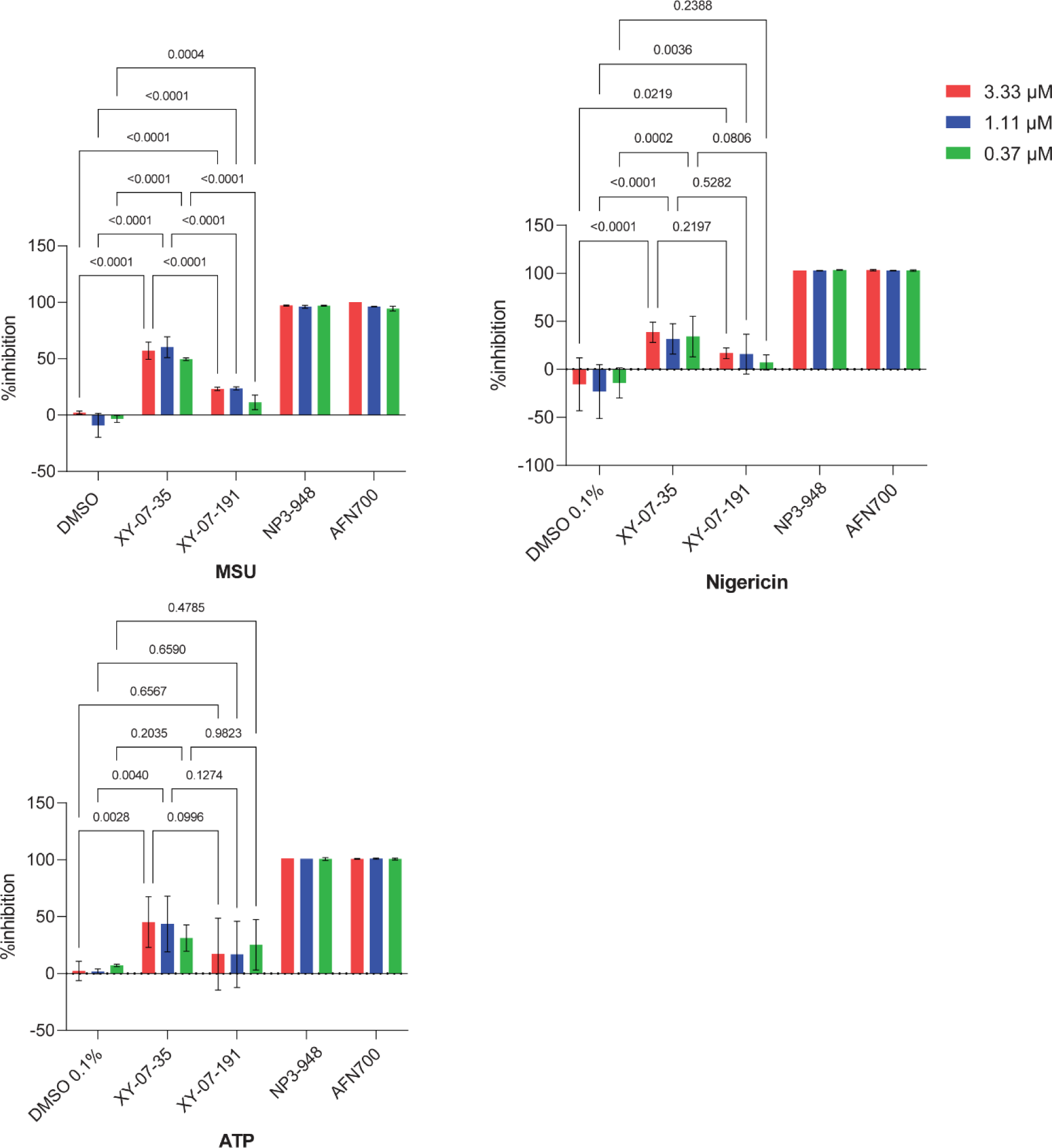
Inhibition of IL-1β release by different treatments (calculated from the data in Fig. 3E). The inhibition obtained with the inflammasome inhibitor NP3-948 at the highest concentration of 3.33 μM represents the maximum inhibition and was set to 100%. The other values were calculated relative to the value of NP3-948, using the following formula: (value condition X – value DMSO) / (value NP3-948 – value DMSO) * 100. Statistical analysis was done by ANOVA.

**Figure S4 (related to Fig. 4):**
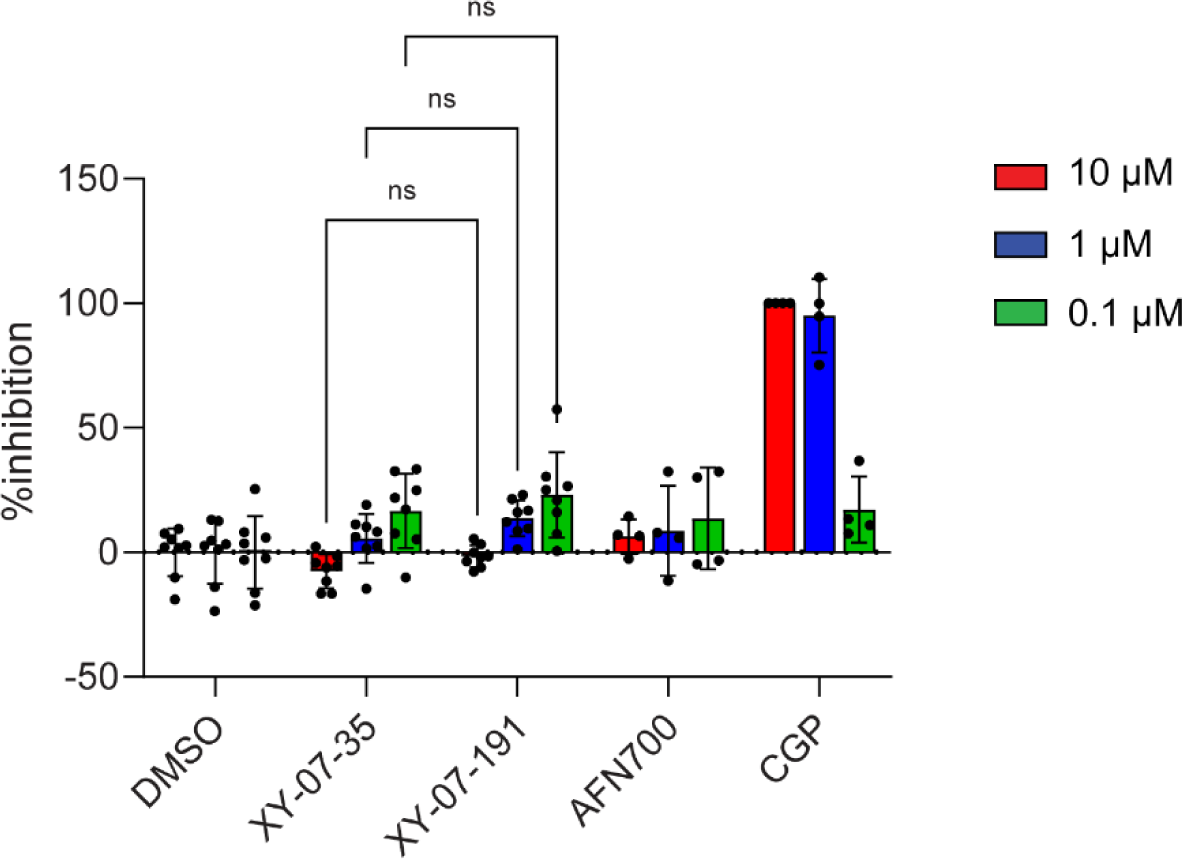
Cell viability was measured by resazurin assay, and the inhibition of cell death is represented as in Fig. S3. The value of CGP treatment at 10 μM was set to 100%.

